# A silent intergenic locus for efficient and versatile transgene expression in *Plasmodium falciparum*

**DOI:** 10.64898/2026.10.13.758768

**Authors:** Laura J. Akkerman, Silvia Tassan-Lugrezin, Ezra T. Bekkering, Freia-Raphaella Lorenz, Geert-Jan van Gemert, Nicholas I. Proellochs, Taco W.A. Kooij, Laura E. de Vries

## Abstract

Genetically modified *Plasmodium falciparum* parasites expressing transgenes such as fluorescent markers are crucial tools for malaria research. However, current reporter lines often rely on integrated transgene constructs that disrupt the coding sequence of a native gene, potentially affecting progression in the life cycle of the tool parasite line. We identified six candidate Silent Intergenic Loci (SILs): putatively editable intergenic regions without transcriptional activity, where gene insertions occurred in *P. falciparum* after its divergence from *Plasmodium berghei* and *Plasmodium vivax*. We characterised the largest candidate SIL on chromosome 12, SIL12, by integration of a constitutively expressed cytosolic mScarlet reporter. The resulting PfRed12 parasites progressed normally throughout the life cycle, from asexual blood-stage replication to mosquito stages and subsequent liver infection. Importantly, they infected both *Anopheles stephensi* and *Anopheles coluzzii* mosquitoes. PfRed12 parasites displayed bright fluorescence in both blood and mosquito stages, making them well suited for live visualisation of parasite transmission. In addition, we used two CRISPR-Cas9 target sites in the mScarlet cassette, one of which we purposely added, to perform efficient re-editing of the transgene. As a proof of concept, we re-edited PfRed12 parasites and integrated an apicoplast metabolic bypass construct, an inducible gametocyte producer construct, and two mNeonGreen reporter constructs. Together, our findings demonstrate that SIL12 permits stable transgene expression without compromising parasite development throughout the life cycle. We established PfRed12 as a new reporter line to study blood- and mosquito-stage development and as a versatile parental line for further genetic engineering.

## 1. Introduction

With more than 250 million cases and over 600,000 deaths annually, malaria remains an immense burden for global health (WHO, 2025). *Plasmodium falciparum* parasites are the causative pathogen of most fatal human malaria cases. *P. falciparum* goes through multiple developmental stages in both the vertebrate and mosquito host, and all stages come with their own set of unique cell biology aspects. Unravelling more about the fundamental biology of these different *P. falciparum* stages is key for the development of novel clinical interventions, such as identifying vaccine and antimalarial targets.

The ability to stably create genetic modifications in the *P. falciparum* genome paved the way for in-depth characterisation of the malaria parasite life cycle (Crabb and Cowman, 1996; Wu et al., 1996). For example, gene modification and deletion facilitated functional analysis (Crabb et al., 1997), and inserting fluorescent markers revealed vital aspect of the parasite’s biology (Kadekoppala et al., 2000). Although generating genetic edited lines in *P. falciparum* remains difficult, the adaptation of CRISPR-Cas9-based protocols has significantly improved the efficiency (Ghorbal et al., 2014), resulting in more complex genetic engineering strategies, and the development of many reporter *P. falciparum* lines (reviewed in (Miyazaki and Miyazaki, 2024)). Examples of reporter lines include parasites that constitutively express (fluorescent) markers to enable efficient quantification and tracking (*e.g.* GFP-NanoLuciferase (Miyazaki et al., 2023)), parasite lines that allow for easy quantification of sexual commitment (*e.g.* NF54-gexp02-Tom (Portugaliza et al., 2019)), or parasites that express fluorophores in specific organelles (*e.g.* PV-mRuby (Glushakova et al., 2018) and MitoRed (Verhoef et al., 2024)). Additionally, multiple tool parasite lines were developed that express transgenes for specific research purposes. These include lines in which sexual commitment can be actively induced to increase gametocyte numbers (*e.g.*iGP2 (Boltryk et al., 2021)), lines that allow for more efficient genetic modifications (*e.g.* NF54::CRISPR (Nasamu et al., 2021), and 3D7^attB^ (Nkrumah et al., 2006)), lines that facilitate conditional deletions (NF54::DiCre (Tibúrcio et al., 2019)), or lines that harbour an alternative metabolic pathway for the production of isopentenyl pyrophosphate to allow for apicoplast disruption (PfMev (Swift et al., 2020)).

There is, however, a lack of established genome safe harbour loci in the *P. falciparum* genome, analogous to *e.g.* the *ROSA26* locus in mice (Friedrich and Soriano, 1991). Therefore, many of the reporter and tool lines in *P. falciparum* have their transgenes placed directly into normally intact genes. Although these genes are thought to have dispensable roles during *in vitro* culture of the asexual blood stages, transgene insertions in some of these sites resulted in abortive phenotypes in other life-cycle stages. For example, *p230p* (Pf3D7_0208900) is now known to have a functional role during gamete fertilisation in the mosquito midgut (Marin-Mogollon et al., 2018), *pfs47* (Pf3D7_1346800) is essential for parasite immune evasion in African *Anopheles gambiae* complex mosquitoes (Molina-Cruz et al., 2013), and the *Plasmodium berghei* orthologue of *lisp1* (Pf3D7_1418100) has been shown to have an essential role during liver merozoite egress (Ishino et al., 2009). Another commonly used genetic locus is *rh3* (PF3D7_1252400), which is annotated as a pseudogene, yet its transcript has been detected in blood stages and its protein product in sporozoites (Taylor et al., 2001; Florens et al., 2002). Finally, *cg6* (Pf3D7_0709200) appears to be dispensable for the whole life cycle (Nkrumah et al., 2006), although it is conceivable that future assays may reveal as yet unknown important functions under specific conditions.

Therefore, in a previous study, we set out to identify neutral genetic loci in the *P. falciparum* genome, using the approach applied successfully for *P. berghei* to identify ‘silent intergenic loci’ (SILs) at chromosomal breakpoints (Kooij et al., 2012). Of three potential sites, we successfully integrated a cassette expressing a mitochondrial-localised fluorescent marker in *P. falciparum* chromosome 7 (Verhoef et al., 2024). Although this line provided robust numbers of both asexual and sexual blood stages that were able to infect *Anopheles stephensi* mosquitoes, the parasites seemed unable to produce salivary gland-infecting sporozoites, preventing the wider use of this locus. Recently, Sekine *et al*. described a second intergenic locus on chromosome 10 where they integrated a reporter cassette. This proved to be a useful approach to generate a fluorescent reporter line that progressed throughout the life cycle (Sekine et al., 2026).

Here, we describe an adapted strategy of our initial search to identify six additional putative SILs in the *P. falciparum* genome in which a gene deletion or insertion occurred since *P. falciparum* diverged from *Plasmodium vivax* and *P. berghei*. We validated SIL12, the largest of these intergenic loci located on chromosome 12, as a neutral editable locus by integrating an *mScarlet* reporter construct. These PfRed12 parasites progressed through the complete life cycle and expressed mScarlet abundantly throughout. Furthermore, the PfRed12 line is highly versatile and allows for efficient insertion of other transgenes in this background through a pair of validated CRISPR-Cas9 targeting regions. The PfRed12 parasite line is thus a robust platform for the generation of new reporter and tool parasite lines that can progress throughout the *P. falciparum* life cycle.

## 2. Materials and Methods

### 2.1 *P. falciparum* culture conditions

*P. falciparum* NF54 parasites (Ponnudurai et al., 1981) and transfectants were cultured in complete medium (RPMI1640 medium supplemented with 25 mM HEPES, 100 mM hypoxanthine, 24 mM sodium bicarbonate, and 10% A^+^ human serum) and 5% O^+^ human Red Blood Cells (RBCs) (Sanquin, The Netherlands). Cultures were maintained at 37 °C in a low oxygen environment (3% O_2_, 4% CO_2_) under shaking conditions.

### 2.2 Gametocyte culture condition and mosquito infections

All gametocytes were cultured in a previously described semi-automatic shaker set-up (Ponnudurai et al., 1986). To induce gametocytogenesis, trophozoite-stage parasite cultures at 1% parasitaemia were cultured in Albumax medium (RPMI1640 with 25 mM HEPES, 25 mM NaHCO_3_, and 0.5% AlbuMAX-II (Gibco, #11021-045)) for 36 h, as previously described (Graumans et al., 2024). Afterwards, gametocytes were cultured with complete medium until maturation. Exflagellation was quantified on day 12 post induction by mixing the culture 1:1 with 200 μM xanthurenic acid in complete RPMI medium. The solution was incubated in a humid chamber at RT for 15 min before counting exflagellation centres using an improved Neubauer chamber.

*An. stephensi* (Nijmegen Sind-Kasur strain (Feldmann and Ponnudurai, 1989)) and *Anopheles coluzzii* (NGousso strain (Klug et al., 2022)) mosquitoes were reared at 30 °C and 27 °C respectively, at 80% relative humidity under a 12 h:12 h light-dark cycle, with *ad libitum* access to 5% glucose water. To assess mosquito infectivity, day 12 mature gametocytes were added to a bloodmeal as described previously (Ponnudurai et al., 1989). Different batches of gametocytes were used to feed the different mosquito species. Blood-fed mosquitoes were maintained at 26 °C and unfed and partially fed mosquitoes were removed. Oocysts were manually counted in mercurochrome-stained dissected midguts on day 6-7 post infection. Salivary glands of infected *An. Stephensi* mosquitoes were dissected on day 14 post infection to assess presence and fluorescence of sporozoites.

### 2.3 Liver-stage culture conditions

Liver-stage parasite culture followed previously described methods (Yang et al., 2021) with minor modifications. Cryopreserved, in-house produced primary human hepatocytes (PHH) from two tissue donors were thawed and seeded onto 96-well culture plates (Corning, #3596) at densities of 0.2 ± 0.02 x10^6^ cells/cm². PHH were maintained at 37 °C and 5% CO_2_ in hepatocyte culture medium, containing William’s Medium E supplemented with GlutaMax (Gibco, #32551-020), 1x insulin/transferrin/selenium (Gibco, #41400-045), 1 mM sodium pyruvate (Gibco, #11360-070), 1x MEM non-essential amino acids solution (Gibco, #11140-035), 1x Penicillin/Streptomycin (Gibco, #15140-122) and 1.6 µM dexamethasone (Sigma-Aldrich, #D4902). Hepatocytes were cultured for two days to allow the formation of a monolayer, before infection with NF54 wildtype or PfRed12 sporozoites. Salivary glands were dissected from infected mosquitoes on day 15 after receiving the gametocyte-enriched bloodmeal and homogenised in hepatocyte culture medium to liberate sporozoites. The suspension was supplemented with 10% human heat-inactivated serum and added onto the hepatocyte monolayer at a sporozoite-to-hepatocyte ratio of 1:1. Plates were centrifuged at 750 g for 5 min and incubated at 37 °C and 5% CO_2_. Culture medium was changed after 4 h, and on day 3 and 5 post infection.

Live fluorescence microscopy was performed on day 5 post infection as described below. Hepatocyte infection rates and parasite growth were assessed on day 3 and 7 post infection on fixed samples. Culture plates were incubated for 10 min in 4% formaldehyde (ThermoFisher, #28906) in PBS and stained overnight at RT with mouse anti-*Pf*GAPDH 7.2 (European Malaria Reagent Repository, supplied by Dr. Jana McBride, 1,000x diluted) (Hall et al., 1983) or rabbit anti-*Pf*HSP70 (Stress Marq, #SPC-186D, 200x diluted) in staining buffer (1% bovine serum albumin (Sigma-Aldrich, #A7906), 0.2% Triton X-100 (Promega, #H5141) in PBS). A secondary anti-mouse Alexa Fluor 647 (Invitrogen, #A-21235, 500x diluted) or anti-rabbit Alexa Fluor 647 antibody (Invitrogen, #A-21245, 500x diluted) was added in staining buffer for 1 h at RT. DNA was stained with 1 µg/ml DAPI solution (Sigma-Aldrich, #MBD0015) in PBS for 30 min at RT. A minimum of 12 non-overlapping images were taken from the central area of each infected well on an Axio Observer 7 Inverted LED microscope with a 10x air objective. Images were analysed in Fiji using an automated thresholding and segmentation pipeline (Schindelin et al., 2012). Extracted features were the total number of segmented nuclei and the number and area of segmented parasites. Correct particle segmentation was manually verified and adjusted if necessary. Hepatocyte infection rates were calculated per well as quotient of the number of segmented parasites over the difference in number of segmented nuclei and number of parasites.

### 2.4 Plasmid generation

Detailed information on the used oligonucleotides can be found in Table S1. All PCR reactions were performed using Primestar^®^ GXL DNA polymerase (Takara Bio), all other cloning enzymes were obtained via New England Biolabs. Plasmid constructs were sequence verified with Plasmidsaurus Whole Plasmid Sequencing (Oxford Nanopore, R10.4.1).

To generate the homology-directed repair (HDR) template for PfRed12 (pRF0779), the 5′ and 3′ homology regions (HRs) for SIL12 were PCR amplified from NF54 genomic DNA and inserted into the pBAT backbone (Kooij et al., 2012) using NgoMIV+EcoRI and AatII+KpnI restriction/ligation cloning, respectively. The mScarlet expression construct containing i) the *Pfh2b* promoter (PF3D7_1105100), ii) an *mScarlet* gene, and iii) a bidirectional 3′UTR (PBANKA_142660) was PCR amplified from an in-house template (Bekkering et al., 2025) and inserted using EcoRI+HindIII. A CRISPR-Cas9 guide target region (AAAGGAGAAGAACTTTTCACcgg), originally targeting *green fluorescent protein,* was selected using CCTop and included in the overhang of the *h2b-*promoter-facing primer (Stemmer et al., 2015).

For the PfGreen12-H2B HDR template, *mScarlet* from pRF0779 was swapped for *mNeonGreen* using NheI+AflII. The *h2b* promoter was then replaced with the *u60S* promoter (PF3D7_1365900) using AgeI+NheI for the PfGreen12-u60S HDR template. The additional CRISPR-Cas9 guide target region was then removed from both plasmids by AgeI+EcoRI restriction followed by Klenow-based blunt-end cloning to yield the final HDR templates.

The SIL12 5′ and 3′ HRs were also inserted into pGGASelect (NEB) to generate pRF0636 to allow for Golden Gate-based cloning. The HA-tagged Super Folder Green apicoplast marker and apicoplast metabolic bypass construct were directly amplified from the genomic DNA of PfMev, a kind gift of Sean T. Prigge (Swift et al., 2020), and then inserted into pRF0636 using the NEBridge^®^ Golden Gate Assembly Kit (BsmBI-v2) to yield the PfMev12 HDR template. Finally, the HDR template of PfiGP12 was created using the same BsmBI-v2 based kit by combining pRF0636 with PCR amplicons of i) the *cam* promoter (PF3D7_1434200), ii) the NF54 *gdv1* coding sequence (PF3D7_0935400), iii) the GlmS ribozyme, and iv) *Pbdhfr* 3’UTR (PBANKA_0719300).

To generate the Cas9 guide plasmids, pairs of complementary oligonucleotides (Table S1) were treated with T4 polynucleotide kinase, annealed, and ligated into BbsI-digested pRF0277. This plasmid was derived from MLB626 (a kind gift from Marcus Lee) (Lim et al., 2016), in which the *hDHFR* cassette was replaced by a *hDHFR-ScFcy1-ScFur1* cassette to facilitate 5’fluorocytosine-based (5’FC) negative selection.

### 2.5 *P. falciparum* genome engineering

For transfections, 60 µg of the Cas9 guide(s) and 60 µg linearised HDR plasmids were ethanol precipitated and dissolved in cytomix (10 mM K_2_HPO_4_/KH_2_PO_4_ pH 7.6, 120 mM KCl, 0.15 mM CaCl_2_, 5 mM MgCl_2_, 25 mM HEPES, 2 mM EDTA) in a total volume of 300 µl. The solution was then added to 400 µl of packed ring-stage infected RBCs and electroporated at 0.31 kV and 950 µF (Wu et al., 1995; Crabb and Cowman, 1996). Parasites were left to recover for 4 h in complete medium and then treated with 2.5 nM WR99210 (Jacobus Pharmaceuticals) for 5 days. To obtain an isogenic population for PfRed12, 250 parasites were isolated by fluorescence-activated cell sorting (FACS) based on the cytosolic mScarlet signal using a CytoFLEX SRT benchtop Cell Sorter (Beckman Coulter). Successful integration was confirmed by diagnostic PCR (Table S1).

Prior to re-transfection, PfRed12 parasites were cultured in the presence of 1 µM 5’FC (Sigma-Aldrich, F7129) for 7 days to remove any left-over episomal SIL12-targeting guide plasmids. Treated parasites were transfected as described above, using different combinations of guide and HDR plasmids. The PfiGP12 parasite was constantly cultured in complete medium supplemented with 2.5 mM glucosamine (GlcN). An isogenic population of 250 parasites was obtained by FACS based on either mNeonGreen signal (PfGreen12-H2B and PfGreen12-u60S) or Super Folder Green signal (PfMev12). To obtain an isogenic population for the marker-free PfiGP12 parasite line, the transfected line was first stained with 0.1 µg/ml Rhodamine123 for 15 min, washed in complete medium, and 250 single cells selected by FACS that were Rhodamine123-positive and mScarlet-negative.

### 2.6 Asexual blood-stage growth assay

Tightly synchronised ring-stage parasites were obtained by a 10-minute treatment with 5% sorbitol (Sigma-Aldrich) in milli-Q, followed by a second sorbitol treatment 16 h later, after which parasitaemia was determined through flow cytometry. For the growth assay, cultures were set up at 0.5% parasitaemia and parasites growth was followed for 4 days. Samples for flow cytometry were taken directly after set-up, at day 1, 2 (before and after dilution), 3, and 4. On day 2, all cultures were diluted with the same dilution factor (1/20). Harvested samples were fixed in 0.25% glutaraldehyde (ITW Reagents, CA0589.100) in PBS and stained with SYBR™ green (1:10,000, ThermoFisher, 7563) for 30 min at 37 °C before analysis with a CytoFlex Flow Cytometer (Beckman Coulter). Data were analysed using FlowJo software (version 10.10).

### 2.7 Live fluorescence microscopy

All microscopy was performed on an Axio Observer 7 Inverted LED microscope equipped with a Colibri 7 LED source, Axiocam 705 mono (Zeiss), and a climate-control incubation housing. Images were taken with either a 10x air objective, 20x air objective, or 63x oil objective using a 385 nm (Hoechst) and 567 nm (mScarlet) LED module, and a brightfield channel. All images were taken on the same day as NF54 wildtype control samples, which were used for identifying the correct visualisation settings in Fiji (Schindelin et al., 2012). For asexual blood stages, gametocytes, and liver stages, PfRed12 and NF54 control parasites were stained with Hoechst33342 (ThermoFisher, H3570, 10,000x diluted) for 15 min at 37 °C and washed into pre-warmed incomplete RPMI1640 medium for blood-stage parasites and hepatocyte culture medium for liver-stage parasites. Parasites were allowed to settle in pre-warmed poly-L-lysin coated 8-well µ-slides (Ibidi), and images were taken at 37 °C. For male and female gametes, a stage V gametocyte culture was incubated in a poly-L-lysin coated µ-slide filled with exflagellation medium (complete medium + 100 μM xanthurenic acid) for 12 min at RT. To compensate for the quick movement of the flagellar microgametes, male (micro)gametes were imaged with a very short exposure time. To image ookinete maturation in the blood bolus, two *An. stephensi* midguts 1 day post infection were crushed in PBS and the diluted contents added to a room-temperature µ-slide. To image midgut oocysts and salivary gland sporozoites, mosquito tissues were dissected on day 9 and day 16 post infection, respectively, and transferred to a PBS-filled µ-slide. To image the thorax of a PfRed12-infected mosquito, an *An. stephensi* mosquito day 16 post infection was doused in 70% ethanol prior to imaging on a PBS-filled µ-slide (Ibidi).

### 2.8 Dose-response assay

Dose-response curves were obtained in a replication assay as previously described (Dery et al., 2015). In short, parasites were incubated with FR-900098 (Sigma-Aldrich, F8307) for 72 h in 96-well plates with or without 50 µM mevalonate, supplied as 50 µM mevalonolactone (Sigma-Aldrich, M4667). Parasites were then lysed in SYBR Green-containing buffer, followed by measurement of fluorescence intensity on a BioTek Synergy 2 plate reader. Signals were normalised to a 0.1% DMSO control (100% growth) and a 1 µM dihydroartemisinin control (0% growth).

### 2.9 Commitment assay

Asexual blood-stage parasites were synchronised by enriching schizonts by MACS using LS columns (Miltenyi Biotec). At 20-24 h post invasion, parasites were set up at 1% parasitaemia in 5% haematocrit in 24-well plates (ThermoFisher, #142475) in complete medium, with or without 2.5 mM GlcN. After 36 h, media was changed for complete RPMI media supplemented with heparin (43 U/ml) to ensure only a single round of sexual commitment. Conversion rates were calculated based on Giemsa-stained smears taken at 36 h (rings) and day 5 (gametocytes) post-induction, counting a minimum of a thousand RBCs.

## 3. Results

### 3.1 Selection of candidate silent intergenic loci (SILs)

To identify candidate SILs, we selected loci in the *P. falciparum* genome where gene insertions or deletions occurred (Kooij et al., 2005; Carlton et al., 2008). Specifically, we focussed on loci that changed after the *P. falciparum* lineage diverged from their common ancestor with *P. vivax* and *P. berghei* (Figure 1A). To find gene insertions, we took the list of *P. falciparum* protein-coding genes in PlasmoDB and excluded all genes with annotated syntenic orthologues in *P. berghei* or *P. vivax* (Alvarez-Jarreta et al., 2024). To find gene deletions, we obtained a list of protein-coding genes in *P. berghei* with an orthologue in *P. vivax* but not in *P. falciparum.* From the resulting lists, we excluded unsuitable sites (Figure 1B), starting with sites where annotation seemed to be incomplete and no insertion or deletion seemed to have occurred. More sites were excluded because of their location within a larger insertion or highly variable region, such as the sub-telomeric regions of a chromosome. One site was excluded because it had previously been tested and plasmid generation by molecular cloning was found to result in frequent spontaneous plasmid rearrangements and very low plasmid yields (Verhoef et al., 2024). We also excluded all intergenic regions containing pseudogenes, since some pseudogenes are expressed and possibly functional (Bethke et al., 2006; Macedo-Silva et al., 2017; Das et al., 2026), or incorrectly annotated. For the remaining loci, we selected sites where the flanking genes were in a tail-to-tail configuration to avoid unwanted editing of (unannotated) promoter regions. Furthermore, we checked for evidence of translational activity within the intergenic regions (Alvarez-Jarreta et al., 2024), and resized candidate SILs to leave these areas intact. Next, we selected sites where there was at least one *Sp*Cas9 guide target sequence with >0.4 on-target efficiency and zero or one relevant off-target hits (Ribeiro et al., 2018). This selection process yielded six candidate SIL sites (Table 1), which are all gene insertion sites.

**Figure 1:**
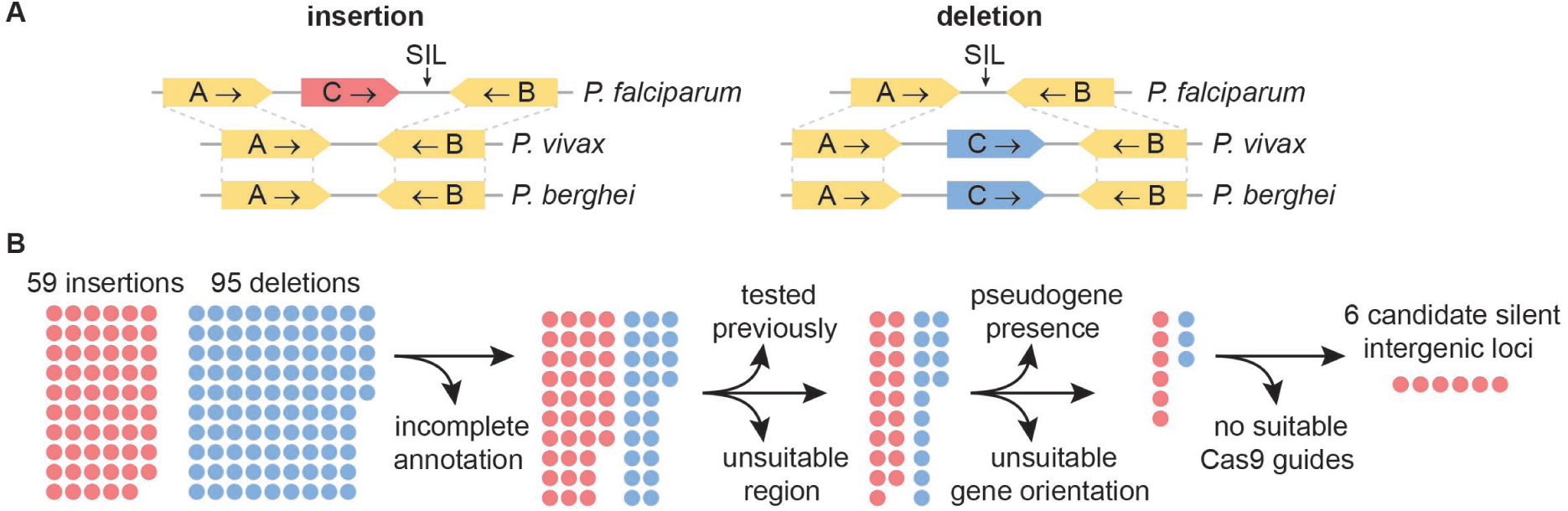
Concept and selection of silent intergenic loci (SILs). A) Schematic representation of insertion and deletion SIL sites. B) Schematic representation of the candidate SIL selection process.

**Table 1:**
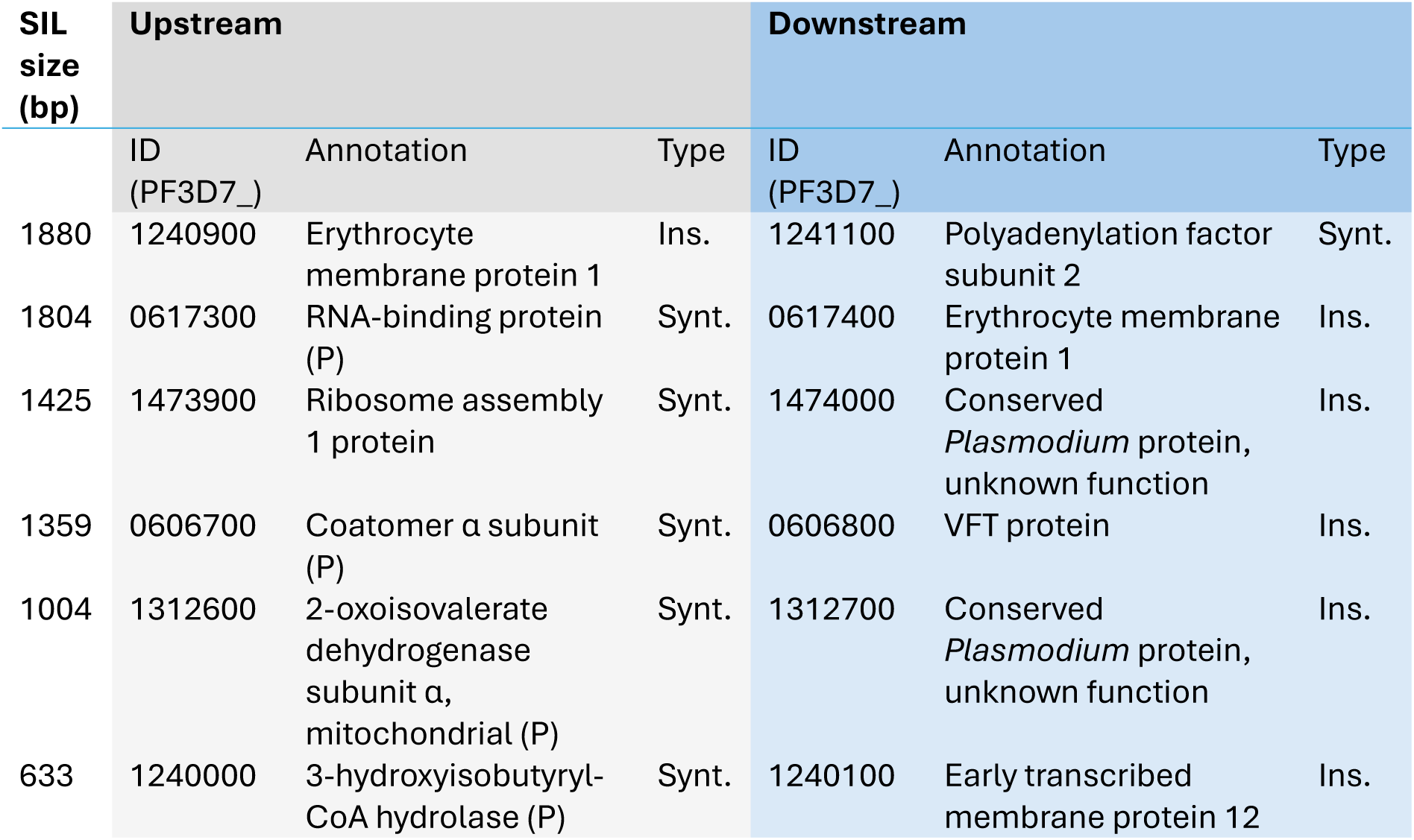
Candidate silent intergenic loci (SILs) at gene insertion sites in the P. falciparum genome. Ins.: P. falciparum-specific insertion. Synt.: syntenic gene. P: putative. Annotations obtained from PlasmoDB (Alvarez-Jarreta et al., 2024).

### 3.2 Validation of SIL12 as a suitable locus for reporter gene expression

We selected the largest candidate SIL, to have the lowest chance of affecting expression of neighbouring genes, for validation. We called the locus SIL12 after its location on chromosome 12. In NF54 parasites, we integrated a gene for cytosolic mScarlet expression under the *Pfh2b* promoter in SIL12 and obtained an isogenic population by FACS (Figure 2A-B). PfRed12 asexual blood-stage parasites replicated normally (Figure 2C) and produced gametocytes when exposed to AlbuMAX-supplemented medium as reported previously (Graumans et al., 2024). Upon a temperature drop and addition of xanthurenic acid, stage V male PfRed12 gametocytes activated and formed exflagellation centres similar to NF54 (Figure 2D), indicating male parasites can undergo gametogenesis. PfRed12 parasites were then fed to mosquitoes and midguts were dissected on day 7 to assess oocyst formation. PfRed12 parasites were able to infect *An. stephensi* (Figure 2E) with a normal oocyst intensity and prevalence, and PfRed12 parasites can also infect *An. coluzzii* (Figure 2F). Furthermore, these parasites were able to progress to salivary gland sporozoites which could successfully infect primary human hepatocytes (Figure 2G, S1). As an indicator of liver-stage development, we measured liver schizont size, which was comparable between NF54 and PfRed12 (Figure 2H).

**Figure 2:**
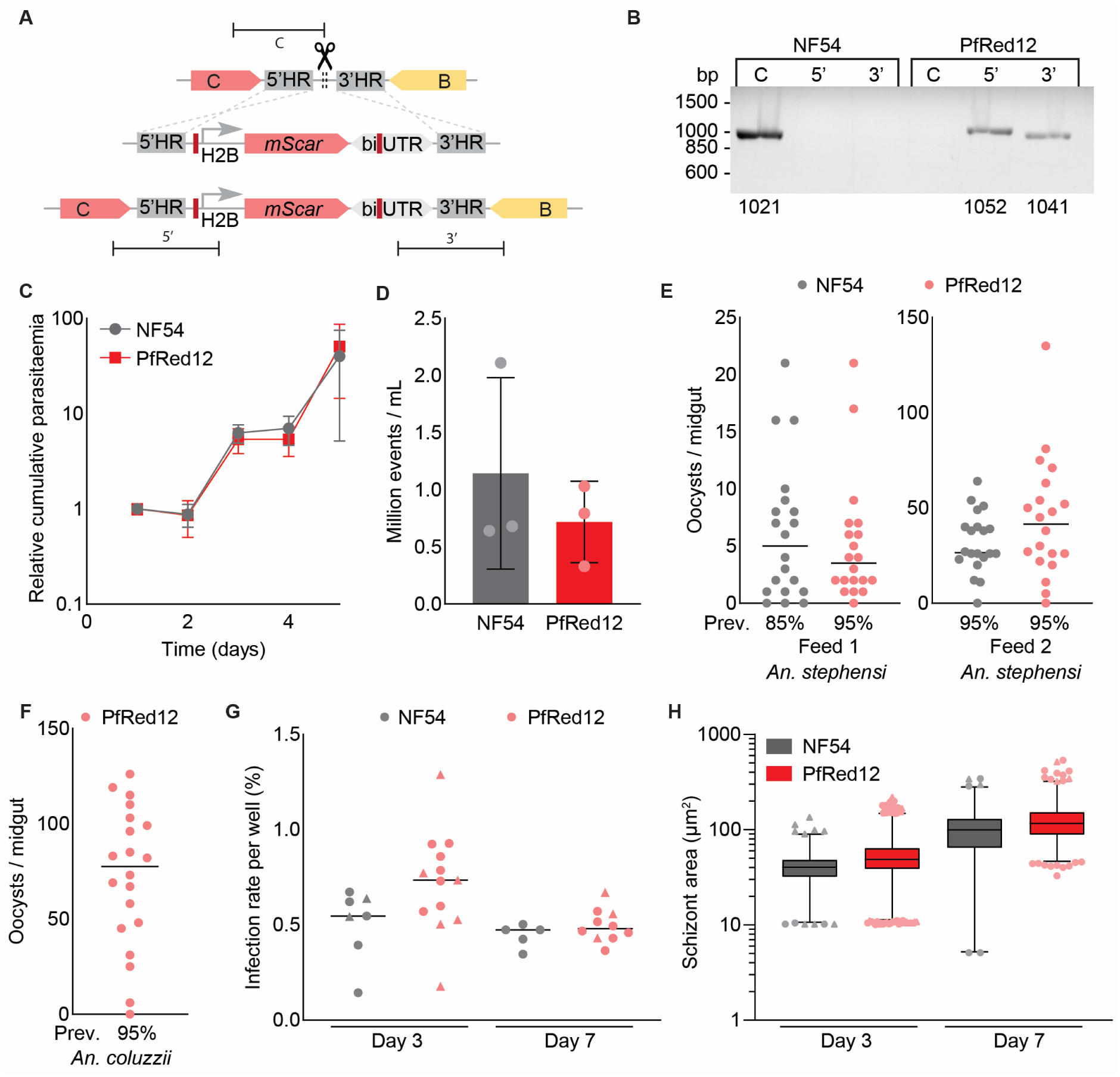
PfRed12 parasites progress normally throughout the life cycle. A) Schematic overview of the strategy to generate PfRed12. The repair plasmid contains a Pfh2b promoter (PF3D7_1105100) to drive expression of mScarlet followed by a bidirectional 3’UTR (PBANKA_142660). Cas9 guide target sequences for re-editing are indicated in dark red. Lines indicate PCR reactions to confirm integration of the construct. B) Diagnostic PCR of PfRed12 with NF54 control to show correct integration and absence of parental line. Expected sizes are indicated below the gel image. C) Relative cumulative parasitaemia of NF54 and PfRed12 asexual blood-stage parasites over time (n = 4). Bars indicate SD. D) Exflagellation of NF54 and PfRed12 gametocyte cultures (n = 3). Bars indicate SD. E) and F) Infection of two times twenty An. stephensi (E) and twenty An. coluzzii (F) mosquitoes with NF54 and PfRed12. Bars indicate median. Prev.: Oocyst prevalence (% of midguts with ≥1 oocyst detected). G) Infection rate per well of primary human hepatocytes infected with NF54 and PfRed12 sporozoites, assessed at day 3 and day 7 post infection. Different symbols indicate different hepatocyte cell donors. Bars indicate median. H) Box & whiskers plot (1-99 percentile) of liver-stage schizont area of NF54 and PfRed12 parasites at day 3 and day 7 post infection of primary human hepatocytes. Different symbols indicate different hepatocyte cell donors.

Using live fluorescence microscopy, we determined whether the reporter gene was successfully expressed. PfRed12 parasites abundantly expressed mScarlet in all stages of the asexual blood-stage replication cycle and during gametocyte development (Figure 3). Not only late-stage asexual blood-stage parasites but also smaller merozoites and ring-stage parasites were easily identified as a result of their bright red colour. PfRed12 parasites also constitutively expressed mScarlet during development in mosquitoes (Figure 4). Male and female gametes, ookinetes, oocysts, and sporozoites were readily observed by live fluorescence microscopy due to abundant expression of the fluorescent marker. Of note, the misalignment between fluorescent and brightfield images of microgametes is due to their continuous rapid movement between the acquisition of different channels during live imaging (Figure 4A). Fluorescent sporozoites were detectable in the mosquito thorax (Figure 4D) and after salivary gland dissection (Figure 4 D-F). Fluorescence of liver-stage schizonts was faintly detectable, although highly auto-fluorescent cell debris that is commonly present in primary human hepatocyte cultures makes visualisation challenging (Figure S2). These observations confirmed that SIL12 can be genetically modified without affecting the parasite’s life cycle under the used conditions and that stable expression of transgenes from the locus is possible in the blood and mosquito stages.

**Figure 3:**
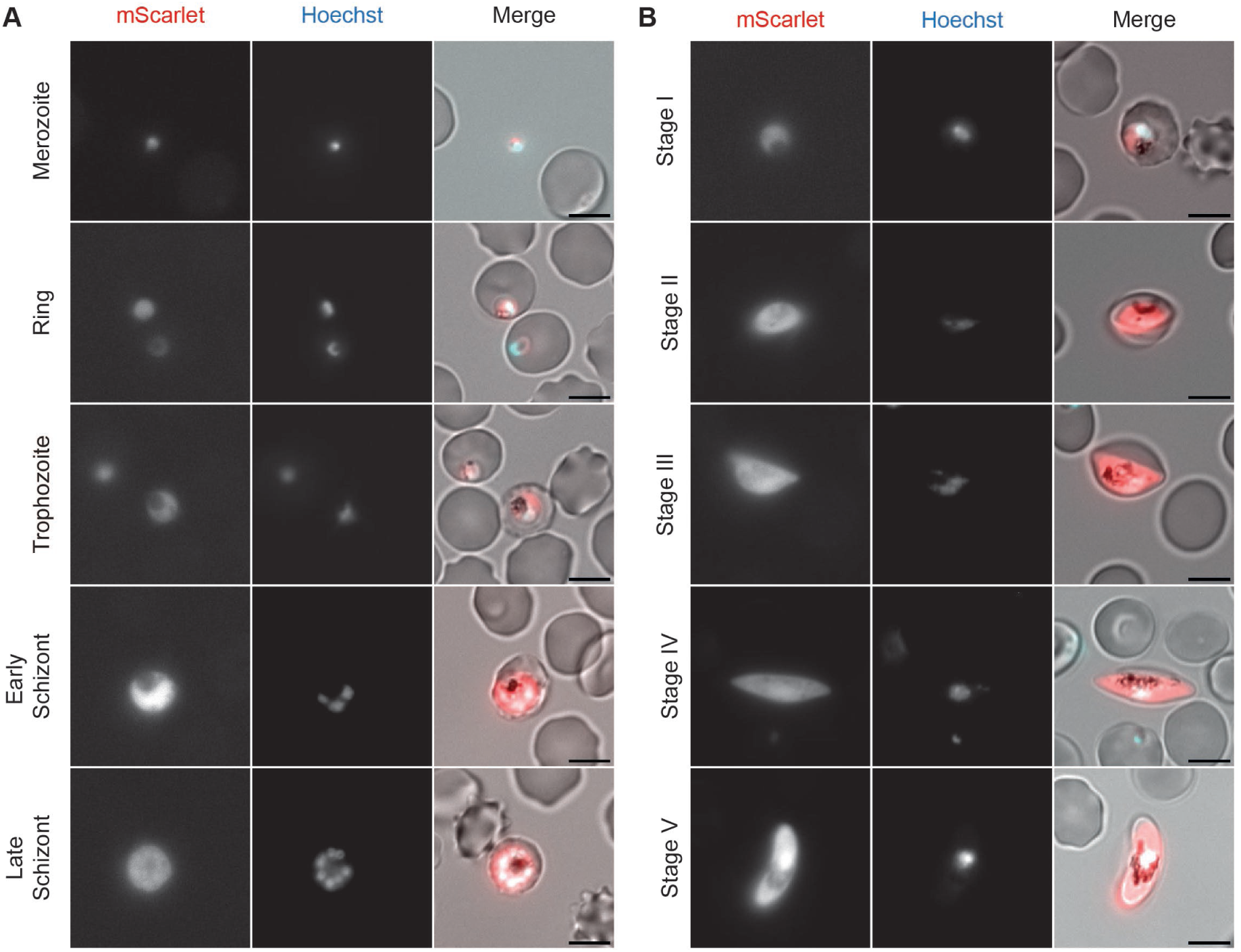
Live fluorescence microscopy for blood-stage PfRed12. Representative images of asexual blood stages (A) and gametocyte stages (B) are shown. Scale bars, 5 µm.

**Figure 4:**
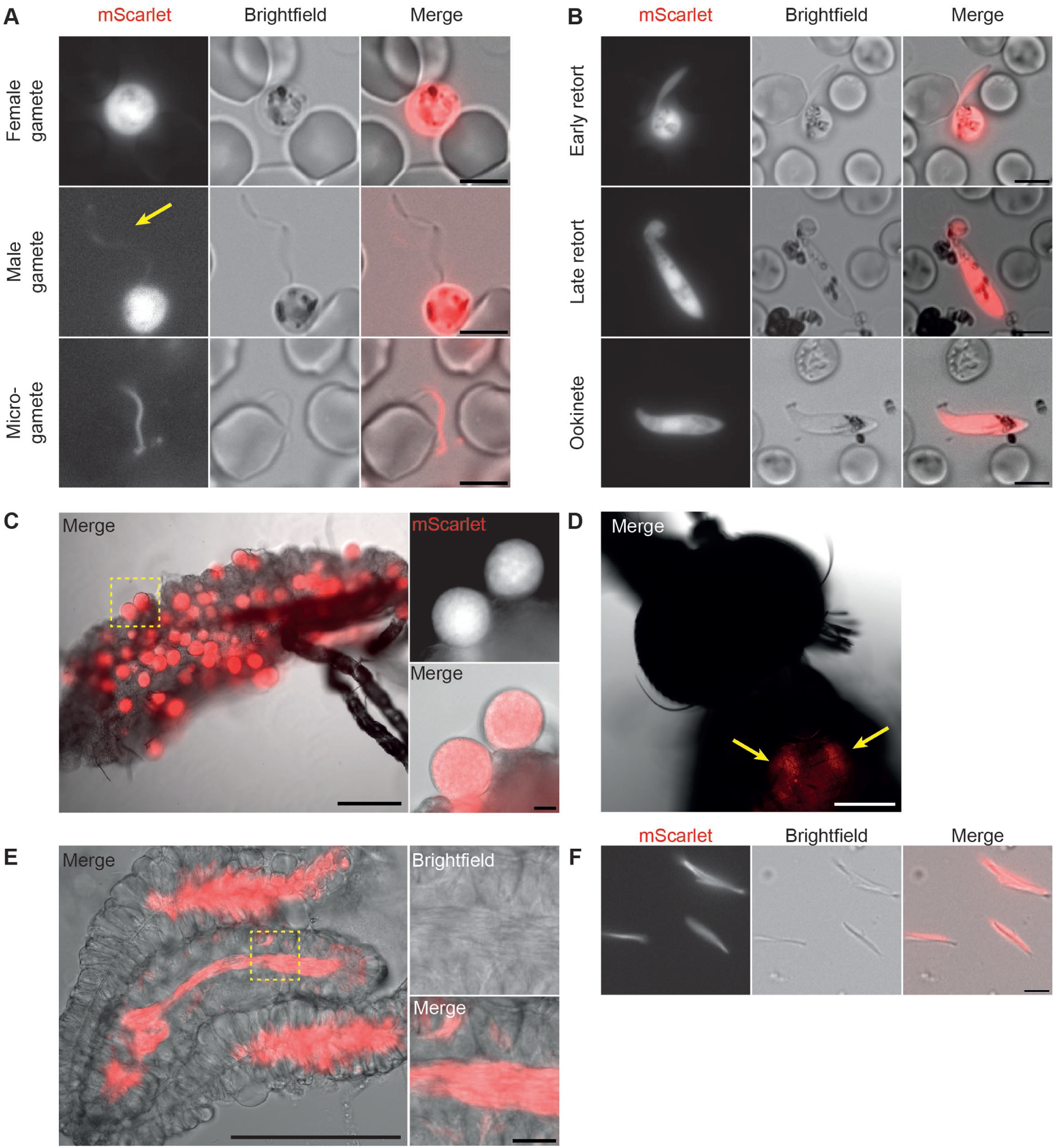
Live fluorescence microscopy of PfRed12 mosquito stages. Representative images of in vitro activated male and female PfRed12 gametes (A), different developmental stages of PfRed12 ookinetes 1 day post infection (B), PfRed12 oocysts from an An. stephensi midgut day 10 post infection (C), a PfRed12 sporozoite-infected mosquito thorax (D) and salivary gland (E), and extracted PfRed12 sporozoites (G). Scale bars, 5 µm (A, B, F), 20 µm (inset C and E), or 200 µm (C-E).

### 3.3 Generation of more tool parasite lines using SIL12 as integration site

The PfRed12 parasite line is a valuable tool as a reporter line but also allows for easy re-editing to integrate other transgenes. During the design of the PfRed12 reporter cassette, we added a unique guide target site between the 5’ homology region and the *h2b* promoter (Figure 2A). Additionally, we selected a Cas9 guide RNA target site in the bidirectional UTR sequence for validation. Both sites have high on-target efficiency scores and no predicted off-target hits in the *P. falciparum* genome (Ribeiro et al., 2018), and we used them to efficiently generate tool parasite lines in the PfRed12 background. In the first line, we swapped the *mScarlet* coding sequence for an *mNeonGreen* coding sequence, obtaining PfGreen12-H2B parasites (Figure 5A-B, S2). In the second line, we exchanged the *h2b* promoter and the *mScarlet* coding sequence for *mNeonGreen* driven by the *u60S* promoter, yielding PfGreen12-u60S parasites (Figure 5C-D, S2). Both lines displayed abundant green fluorescence, with an expected stronger signal when *mNeonGreen* expression was driven by the *h2b* promoter (Figure 5B and 5D). Third, we generated PfMev12 by integrating a metabolic apicoplast bypass and a fluorescent apicoplast marker construct (Figure 5E), which was previously integrated in NF54^attB^ parasites to obtain PfMev (Swift et al., 2020). Just like PfMev parasites (Swift et al., 2020), PfMev12 parasites expressed a fluorescent apicoplast marker and could survive treatment with the apicoplast-targeting compound FR-900098 in the presence of mevalonate (Figure 5F-G, S3).

**Figure 5:**
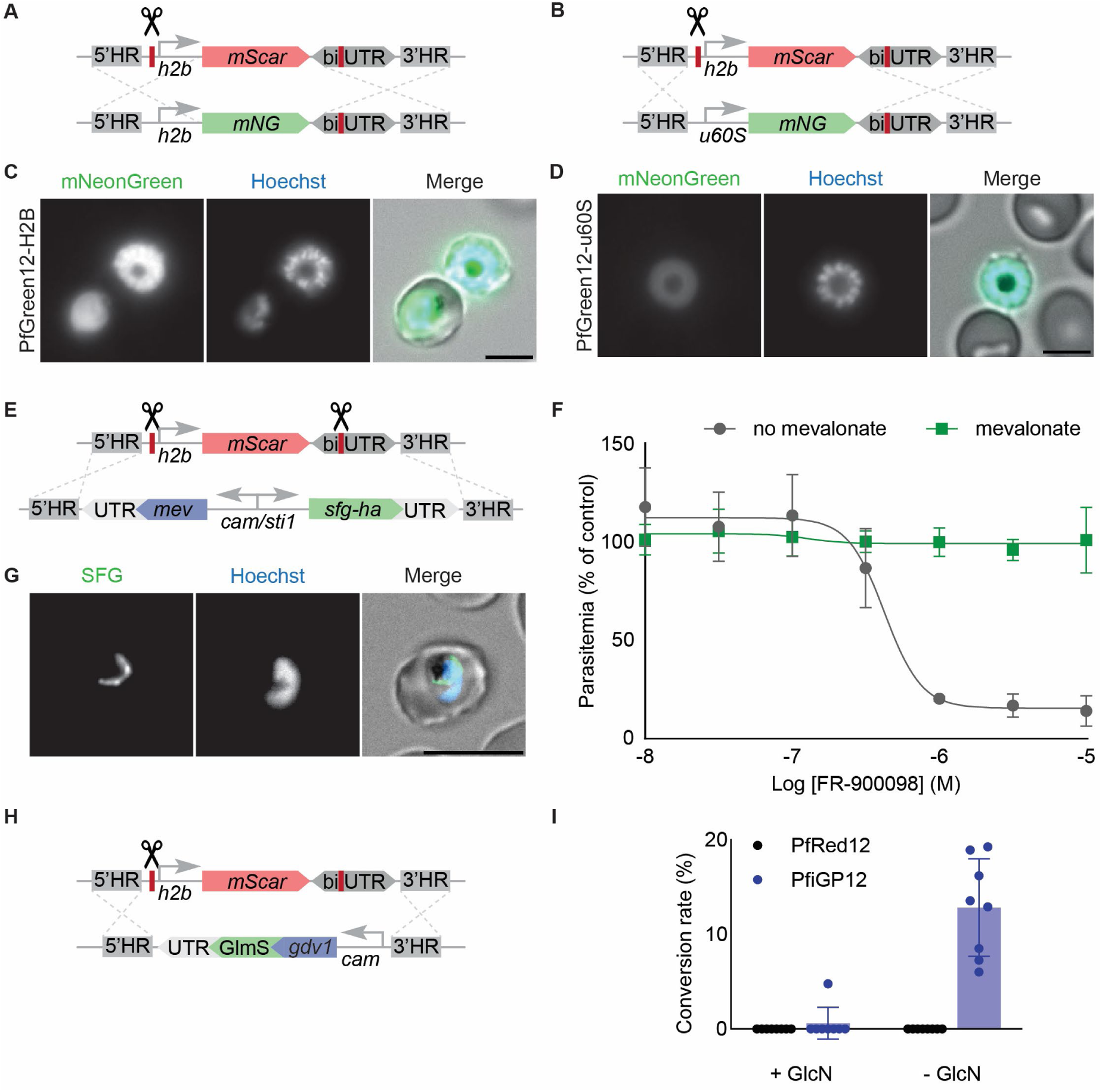
Integration of various tool constructs in SIL12 of PfRed12 parasites. A) Design of PfGreen12-H2B construct. B) Design of PfGreen12-u60S construct. C) Live fluorescence microscopy of PfGreen12-H2B blood-stage schizont. D) Live fluorescence microscopy of PfGreen12-u60S blood-stage schizont. E) Design of PfMev12 construct. F) Dose-response curves *of asexual blood-stage PfMev12 parasites for FR-900098 with or without the presence of 50 µM mevalonate (n = 3). Bars indicate SD. G) Live fluorescence microscopy of PfMev12 blood-stage parasite. H) Design of PfiGP12 construct. I) Commitment assay of PfiGP12 parasites. Bars indicate SD*.

Acquiring a clean marker-free population usually requires obtaining single cell-derived clones, *e.g.* through limiting dilution. This can result in a defect in transmission that is unrelated to the genetic modification, requiring the screening of multiple clones to draw reliable conclusions (McLean et al., 2019; van de Vegte-Bolmer et al., 2021). The PfRed12 tool parasite line allows for obtaining a clean, marker-free, isogenic population by selectively sorting for mScarlet-negative parasites by FACS (Figure S4). Using this approach, we generated PfiGP12, which harbours an inducible *gdv1* gene to drive gametocyte production (Figure 5H-I, S3), a design similar to the iGP2 tool line (Boltryk et al., 2021). Taking all transfections together, we experienced a 71% success rate with edited parasites coming up between 2-3 weeks after transfection, indicating that the used guides are efficient to edit or re-edit SIL12. Together, these results exemplify the versatility of PfRed12 parasites as a tool to generate parasites with a wide variety of applications.

## 4. Discussion

The characterisation of the malaria parasite *P. falciparum* is hampered by the limited availability of loci where transgenes can be integrated efficiently and without affecting life cycle progression. Here, we show SIL12 as a genetically editable intergenic locus in the *P. falciparum* genome by integrating a cytosolic mScarlet marker cassette that facilitates live imaging. These PfRed12 parasites progress normally throughout the life cycle and can infect mosquitoes including those of the *An. gambiae* complex, unlike parasite lines that used *pfs47* as integration locus (Molina-Cruz et al., 2013). Finally, we present PfRed12 as a versatile platform that allows for efficient re-editing, which we demonstrated by generating four additional tool lines.

Most of the currently available *P. falciparum* tool parasite lines have transgenes that are directly integrated into genes that are dispensable in the asexual blood stage. However, many of these gene disruptions are known to cause phenotypes in other stages of the life cycle (Ishino et al., 2009; Molina-Cruz et al., 2013; Marin-Mogollon et al., 2018). Therefore, we aimed to integrate a transgene between two genes, an approach that was indeed successfully trialled in *P. falciparum* by Sekine *et al*. (Sekine et al., 2026) during the preparation of this manuscript. Many intergenic regions, however, are functionally important, for *e.g.* gene regulation or the expression of non-coding RNAs. It can be argued that sites where chromosomal rearrangements, gene insertions, or gene deletions took place are more likely to tolerate further modifications. We previously identified chromosomal breakpoints in *P. berghei* and *P. falciparum* to integrate transgenes without affecting the life cycle, which was successful in the rodent malaria parasite model but less so in *P. falciparum* (Kooij et al., 2012; Verhoef et al., 2024). In order to identify a true safe harbour in *P. falciparum*, we focussed on sites where gene insertions and deletions took place after *P. falciparum* diverged from *P. vivax* and *P. berghei* and we identified SIL12 as a suitable locus to express reporter genes.

SIL12 is an intergenic region on chromosome 12 between (putative) protein-coding genes Pf3D7_1240900 (a *var* gene) and Pf3D7_1241100 (a putative gene for polyadenylation factor subunit 2) (Su et al., 1995). This region contains non-protein-coding gene Pf3D7_1241000, annotated as *Plasmodium* RNA of unknown function 6 (*ruf6*). Overexpression of this GC-rich non-coding RNA gene was reported to disrupt *var* gene expression (Guizetti et al., 2016). Therefore, we avoided disrupting any region with transcriptional activity in SIL12, which leaves 1880 base pairs between the 3’ end of the *var* gene and the 3’ end of *ruf6*. Although SIL12 is located on the 3’ end of two genes in a region with no transcriptional activity, its location might make it less suitable for researchers studying *var* genes. Out of the other five candidate SILs that we identified, one other site also neighbours a *var* gene (Pf3D7_0617400), while the others have neighbouring genes with various (putative) functions and varying (predicted) importance in the asexual blood stage. With SIL12, the intergenic region presented by Sekine *et al*., and the five candidate SILs, the research community now has an expanded toolkit of integration sites. The availability of multiple complementary loci will facilitate the development of complex tool parasite lines, for example with a gene encoding Cas9 or Cre recombinase in one site, and a reporter gene in another.

We envision that PfRed12 parasites can be used in a wide range of experiments that require stable expression of a fluorophore, including but not limited to competition growth assays, genetic crosses, and sporozoite tracking (Münter et al., 2009; Li et al., 2026; Verhoef et al., 2026). PfRed12 was made using a CRISPR-Cas9 guide plasmid that contains both a positive and a negative selection cassette, allowing for easy re-editing using the same positive selection cassette in the next transfection. The mScarlet cassette in PfRed12 can be edited or replaced using various CRISPR-Cas9 guide target sequences, one of which we specifically added. Complete marker-free integration of transgenes is also possible, since the red mother line can be removed through fluorescent cell sorting. As a proof-of-concept that PfRed12 is readily editable to generate (marker-free) tool lines, we generated PfGreen12-H2B, PfGreen12-u60S, PfMev12, and PfiGP12. Importantly, all guides used to edit or re-edit SIL12 have a high on-target efficiency and we achieved a high transfection success rate (Ribeiro et al., 2018). Together, we present SIL12 and PfRed12 as valuable tools for future investigations of the malaria parasite throughout the life cycle.

## CRediT authorship contribution statement

**Laura. J. Akkerman:** Conceptualisation, Investigation, Writing – Original Draft, Funding acquisition; **Silvia Tassan-Lugrezin:** Conceptualisation, Investigation, Writing – Original Draft; **Ezra T. Bekkering**: Conceptualisation, Methodology, Investigation, Writing – Original Draft, Funding acquisition; **Freia-Raphaella Lorenz**: Investigation; **Geert-Jan van Gemert**: Investigation; **Nicholas I. Proellochs**: Conceptualisation, Methodology, Investigation; **Taco W.A. Kooij**: Conceptualisation, Supervision, Funding acquisition, Writing – Original Draft; **Laura E. de Vries:** Conceptualisation, Supervision, Funding acquisition, Writing – Original Draft. All authors reviewed and edited the final draft.

## Acknowledgements

We would like to thank Julie Verhoef for laying the groundwork in search of SILs in *P. falciparum*. We are grateful to Sean Prigge for providing PfMev parasites from which we obtained the apicoplast marker/metabolic bypass construct. We appreciate the help of Aleksandra Nenova, Mariangela Longobardi, and Ghazal Sodaifi with generating some of the plasmids and parasite lines. We thank Rianne Stoter, Laura Pelser-Posthumus, Jolanda Klaassen, and Astrid Pouwelsen for their contributions to the mosquito infection experiments. We also acknowledge the Radboudumc Technology Centre of Flow cytometry for the use of their machines. We thank Stéphanie Blandin for sharing the *An. coluzzii* mosquitoes that were originally colonised by IRD/OCEAC in Yaounde, Cameroon. The anti-*Pf*GAPDH monoclonal antibody 7.2 was obtained from the European Malaria Reagent Repository (http://www.malariaresearch.eu). We thank Hans de Wilt and the Radboudumc Department of Surgery for supplying liver tissue samples, as well as Benjamin Mordmüller and Matthew McCall for supervising the hepatocyte infection assays.

