## Supplementary Material for "A silent intergenic locus for efficient and versatile transgene expression in *Plasmodium falciparum*"

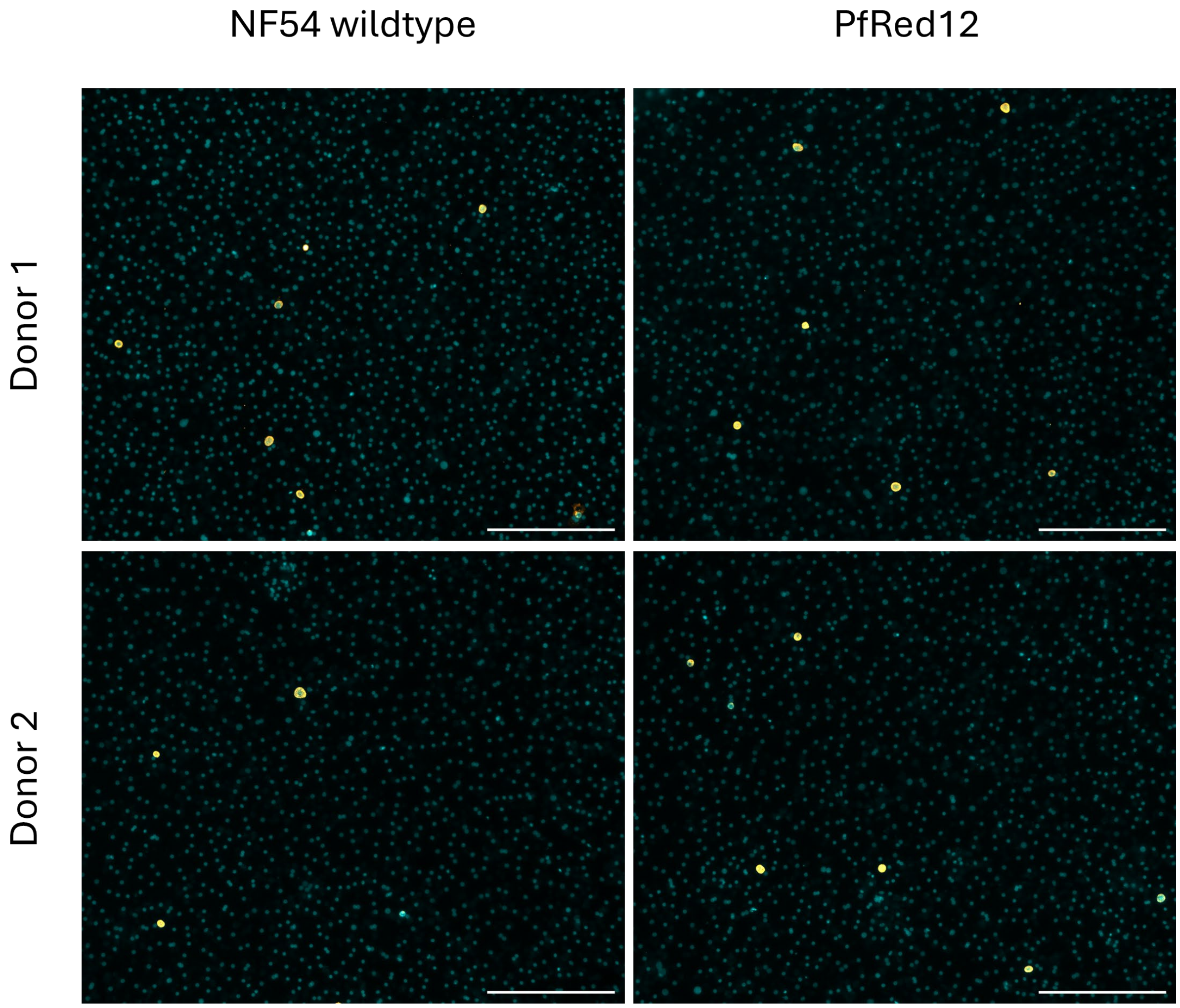


***Supplementary Figure 1: Fluorescence microscopy of liver stage NF54 wildtype and PfRed12.*** *Representative images of liver-stage schizonts on day 7 post infection in two primary human hepatocyte donors. Staining with DAPI (cyan) and anti-Pf-GAPDH (orange). Scale bar 200 µm.*

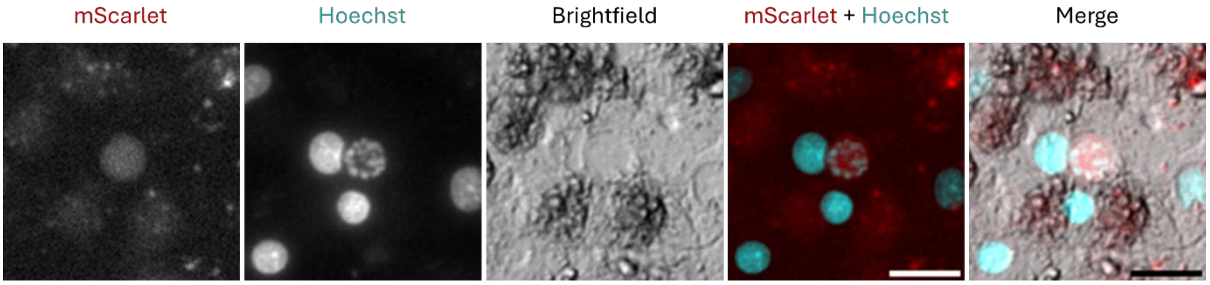


***Supplementary Figure 2: Live fluorescence microscopy of liver stage PfRed12.*** *Representative image of liver-stage schizont on day 5 post infection. Scale bar 10 µm.*

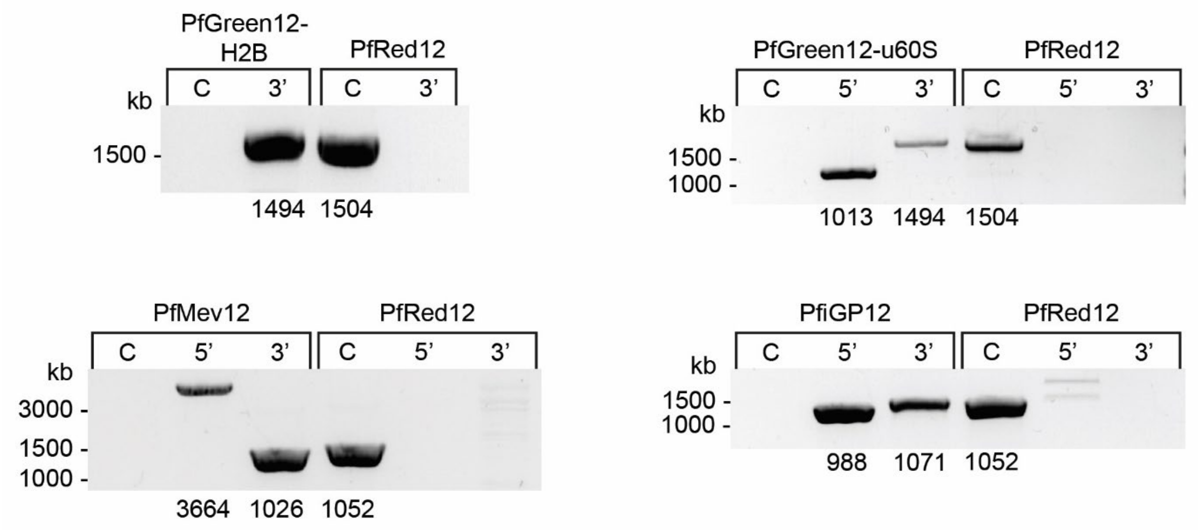


*__Supplementary Figure 3: Diagnostic PCRs of tool parasite lines.__ PCR reactions to check 5' and 3' integration and a control (C) reaction to check the presence of parental parasites were performed for each line. The same reactions were performed on the parental line, PfRed12, as a control. For PfGreen12-H2B, the 5' reaction was omitted because the 5' region (the* h2b *promoter) was present in both the parental line and the edited line.*

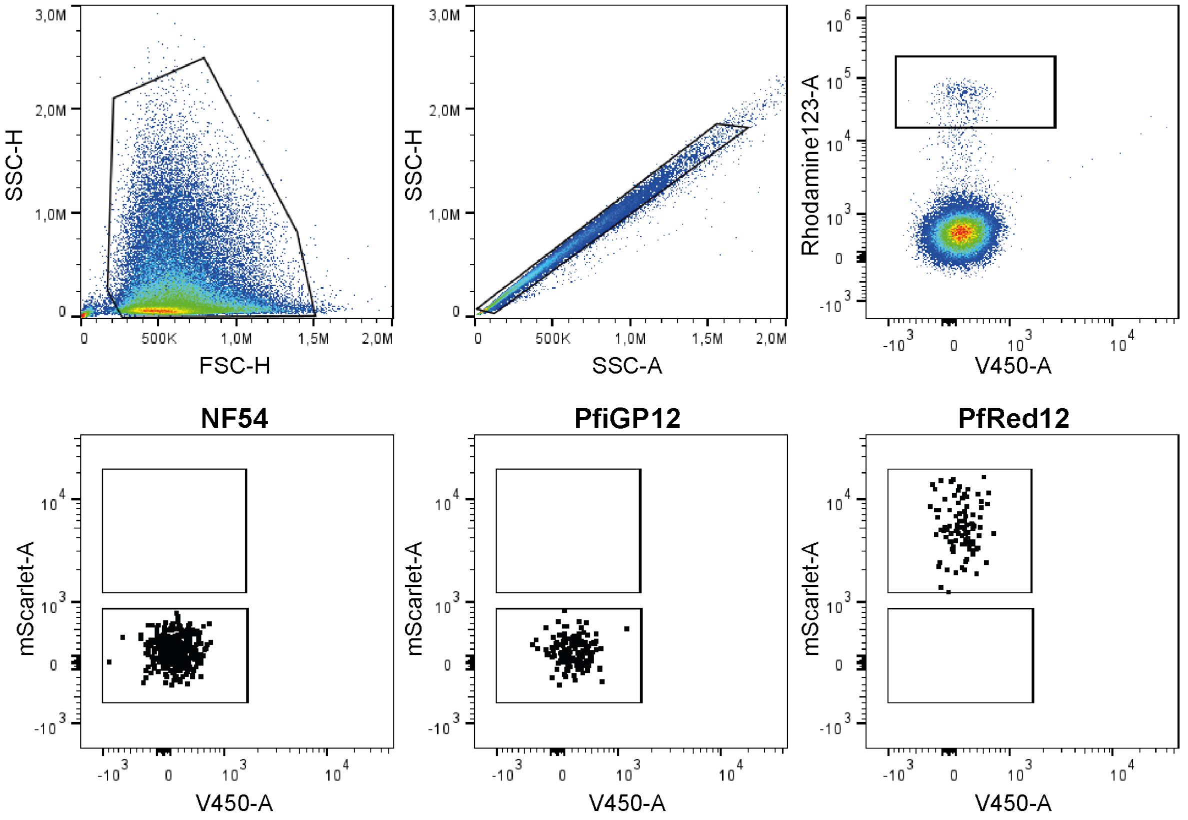


***Supplementary Figure 4: Example of Fluorescence Activated Cell Sorting gating strategy.*** First cellular debris and doublets are excluded based on SSC/FSC signal. Events with high Rhodamine123 staining (active mitochondrial potential) are included. The gating for mScarlet-negative events is based on NF54 (left) and PfRed12 (right) controls.

***Supplementary Table 1: Oligonucleotides list.*** *List of all primers and guides used during this study. Blue nucleotides indicate restriction sites used for cloning.*

| | Function | Sequence |
|---|---|---|
| | HDR template primers | |
| p01 | 5' HR PfRed12 F | AATTTAGCCGGCTTATTTGTCGCTTTTTTGTTTGTGG |
| p02 | 5' HR PfRed12 R | ATTTAAGAATTCAAAACAATTATATTATATTCAGTTATAATGGAG |
| p03 | 3' HR PfRed12 F | AATAAAGGTACCATGATTGAACCAAATGAATAAAAAGGC |
| p04 | 3' HR PfRed12 R | AATAATGACGTCCCCAATCGTTAGGTATGTTGCC |
| p05 | H2B_mSc_biUTR F | CAGTGAATTCGGTCTCTGTGCAAAGGAGAAGAACTTTTCACCGGTTCGAATCAA CTCCGTTCAG |
| p06 | H2B_mSc_biUTR R | CAGTAAGCTTGGTCTCTGACACTAAATGGATGATTCCCCTCTTTGC |
| p07 | mNG F | CAGTGCTAGCATGGTGAGCAAGGGCGAGG |
| p08 | mNG R | CAGTCTTAAGTTACTTGTACAGCTCGTCCATGC |
| p09 | u60S F | TTATACCGGTTAAAATTATTTTCAAAAAAATTGTATGAATATAT |
| p10 | u60S R | CAGTAGCTAGCCATTTTCTTAAAAATAATGAAATG |
| p11 | 5'HR in pRF636 F | GGATCCGGTCTCTGGAGGTCCCCAATCGTTAGGTATGTTGCCTTTC |
| p12 | 5'HR in pRF636 R | AGATCTGGTCTCTTCGTTGTCGTCTCTTGTCCCATGATTGAACCAAATGAATAAA AAGGCAATTTC |
| p13 | 3'HR in pRF636 F | GATCTGGTCTCTACGATCTCGTCTCTGCACCAAAACAATTATATTATATTCAGTTA TAATGGAGACATTG |
| p14 | 3'HR in pRF636 R | ACGTCGGTCTCTATGGCTTATTTGTCGCTTTTTTGTTTGTGGTATTC |
| p15 | PfiGP12 cam promoter F | CAGTTCGTCTCTTCATTGATATATTTCTATTAGGTATTTATTATTATAAAATATAAATC TTG |
| p16 | PfiGP12 cam promoter R | CACTAAGCTTGCGTCTCTGACACAAGAGAAAAGGCAGAAACAAAATAAATTATA ATAAAAAAC |
| p17 | PfiGP12 gdv1 F | CATGTCGTCTCGATGAACTACAAAAAAGAGACTTGCGTAAAATTG |
| p18 | PfiGP12 gdv1 R | CATGTCGTCTCACAGCTTATTTATATGTACATTTTTCTTTTATATAATAATGAATTGG |
| p19 | PfiGP12 GlmS F | CATGAAGCTTGCGTCTCTGCTGTAATTATAGCGCCCGAACTAAG |
| p20 | PfiGP12 GlmS R | CAGTTGGATCCTCGTCTCTCTTACGAGATCATGTGATTTCTCTTTGTTCAAG |
| p21 | PfiGP12 pbdhfr F | CACCAAGCTTGCGTCTCGTAAGGGATATGGCAGCTTAATGTTCGTTTTTC |
| p22 | PfiGP12 pbdhfr F | CAGTTGGATCCTCGTCTCTGTGCCTACCCTGAAGAAGAAAAGTCCGATG |
| p23 | PfMev12 PfMev F | GACTCGTCTCTGACAGAATTCCCAGCATGCTATTTGATGAATTAACTACAC |
| p24 | PfMev12 pfMEVR | CTAGCGTCTCTGTGCGGTCTCTAGCTCGCCAGCCTAGATTTAATAAATATGTTC |
| | CRISPR-Cas9 guide plasmid oligonucleotides | |
| p25 | SIL12 locus guide1 F | TATTGATACAAAATTTTCTAAAAGG |
| p26 | SIL12 locus guide1 R | AAACCCTTTTAGAAAATTTTGTATC |
| p27 | SIL12 locus guide2 F | TATTGTATAAGTGTTCTATAATCAA |
| p28 | SIL12 locus guide2 R | AAACTTGATTATAGAACACTTATAC |
| p29 | Left guide (targets GFP) F | TATTGAAAGGAGAAGAACTTTTCAC |
| p30 | Left guide (targets GFP) R | AAACGTGAAAAGTTCTTCTCCTTTC |
| p33 | biUTR guide F | TATTGCTTCAATGGATAATGTATAG |
| p34 | biUTR guide R | AAACCTATACATTATCCATTGAAGC |
| | Diagnostic PCR to check genomic integration | |
| p35 | PfRed12 - C F | CTAAAAGGAGGTTCTACAAGAAACC |
| p36 | PfRed12 - C R | CTTTTATATTAGTACTAATATTAAGCTGCC |
| p37 | PfRed12 - 5'HR F | GCACATCCTTATATACATTATGCACATTTATGTATG |
| p38 | PfRed12 - 5'HR R | p36 |
| p39 | PfRed12 – 3'HR F | TGTGTGTCTATATATTTTGTATGTTTGC |
| p40 | PfRed12 – 3'HR R | CGATTTCTTAATATTGATAAGTATCATGTG |
| p41 | PfGreen12-H2B - C F | GGTGGAACAGTACGAACGCTCC |
| p42 | PfGreen12-H2B - C R | p36 |
| p43 | PfGreen12-H2B - 3'HR F | CTCAACTTCAAGGAGTGGCAAAAG |
| p44 | PfGreen12-H2B - 3'HR R | p36 |
| p45 | PfGreen12-u60S - C F | p41 |

| | | |
|---|---|---|
| p46 | PfGreen12-u60S - C R | p36 |
| p47 | PfGreen12-u60S – 5'HR F | p39 |
| p48 | PfGreen12-u60S – 5'HR R | CATTTTTCTTTACATAAAAAAACTGTCCAATAAAAAATG |
| p49 | PfGreen12-u60S - 3'HR F | p43 |
| p50 | PfGreen12-u60S - 3'HR R | p36 |
| p51 | PfiGP12 - C F | p37 |
| p52 | PfiGP12 - C R | p36 |
| p53 | PfiGP12 – 5'HR F | p39 |
| p54 | PfiGP12 – 5'HR R | GACAGCTCAATTCTTTATGTCCACAAC |
| p55 | PfiGP12 - 3'HR F | GCATAACAAAAAATATATTCATACAAATATAAAAAAATGTG |
| p56 | PfiGP12 - 3'HR R | p36 |
| p57 | PfMev12 - WT F | p37 |
| p58 | PfMev12 - WT R | p36 |
| p59 | PfMev12 - 5'HR F | p39 |
| p60 | PfMev12 - 5'HR R | CCGCGTTTAGTTTTGTTATTTAGC |
| p61 | PfMev12 - 3'HR F | GTGTAGTTAATTCATCAAATAGCATGCTGG |
| p62 | PfMev12 - 3'HR R | p36 |
